# Fast and exact gap-affine partial order alignment with POASTA

**DOI:** 10.1101/2024.05.23.595521

**Authors:** Lucas R. van Dijk, Abigail L. Manson, Ashlee M. Earl, Kiran V Garimella, Thomas Abeel

## Abstract

**Motivation:** Partial order alignment is a widely used method for computing multiple sequence alignments, with applications in genome assembly and pangenomics, among many others. Current algorithms to compute the optimal, gap-affine partial order alignment do not scale well to larger graphs and sequences. While heuristic approaches exist, they do not guarantee optimal alignment and sacrifice alignment accuracy.

**Results:** We present POASTA, a new optimal algorithm for partial order alignment that exploits long stretches of matching sequence between the graph and a query. We benchmarked POASTA against the state-of-the-art on several diverse bacterial gene datasets and demonstrated an average speed-up of 4.1x and up to 9.8x, using less memory. POASTA’s memory scaling characteristics enabled the construction of much larger POA graphs than previously possible, as demonstrated by megabase-length alignments of 342 *Mycobacterium tuberculosis* sequences.

**Availability and implementation:** POASTA is available on Github at https://github.com/broadinstitute/poasta.

## Introduction

Multiple sequence alignments (MSAs) are central to computational biology. MSAs have many applications, including computing genetic distances, which can serve as a basis for a phylogeny; determining consensus sequences, e.g., to perform read error correction; and identifying allele frequencies, e.g., for sequence motif identification.

Computing the optimal MSA with the *sum of all pairs* (SP) score is an NP-complete problem (Wang and Jiang, 1994). These classical exact algorithms have a runtime exponentially related to the number of sequences and are thus intractable for even modest-sized datasets. Instead, nearly all popular MSA tools, including MAFFT (Katoh and Standley, 2013) and MUSCLE (Edgar, 2004), compute the MSA progressively: first, an alignment between two sequences is computed, then additional sequences are added one by one until all sequences have been aligned. The runtime of these approaches is linear in the number of sequences instead of exponential. While MSAs computed this way do not necessarily find the globally optimal solution for the SP objective, they are still highly useful approximations to otherwise intractable alignment problems.

Partial order alignment (POA) is a well-known progressive MSA approach that pioneered using a graph to represent an MSA rather than a sequence profile (Lee et al., 2002). This improved the ability to represent indels, leading to higher-quality alignments. Since POA is a progressive MSA algorithm, the optimal SP score is not guaranteed for the entire MSA. However, POA does guarantee that each individual sequence-to-graph alignment is optimal.

POA is relevant to many applications, including *de novo* genome assembly (e.g., read error correction and consensus generation) (Chin et al., 2013; Loman et al., 2015; Vaser et al., 2017), RNA isoform inference (Lee, 2003), structural variant (SV) characterization (Chaisson et al., 2019), and variant phasing (Holt et al., 2023).

POA is also essential to two recent human pangenome graph construction pipelines (Hickey et al., 2023; Garrison et al., 2023). These pipelines are pushing the limits of POA, as aligning long stretches of homologous sequence among input genomes requires substantial computing and memory resources. For example, consider the gap-affine alignment of a 500 kbp sequence to a graph with 500k character-labeled nodes. Conventional POA approaches have a runtime and memory complexity of *O*(*Nm*), i.e., a product of the POA graph size *N* and the sequence length *m*. This example would, therefore, require about 3 TB of RAM (assuming 32-bit integers for storing alignment costs in three alignment state matrices).

Several tools, including SPOA (Vaser et al., 2017) and abPOA (Gao et al., 2021), have been developed to address the need for faster and more memory-efficient POA algorithms. The current state-of-the-art, SPOA, is a reimplementation of the original algorithm, which accelerates computing the dynamic programming (DP) matrix by using single-instruction-multiple-data (SIMD) instructions available on modern CPUs. While faster, SPOA still computes the full DP matrix and thus does not ameliorate demands on memory usage. abPOA additionally improves performance and memory usage by applying an adaptive banding strategy to partially compute the DP matrix. However, this sacrifices the guarantee of finding the optimal sequence-to-graph alignment.

Here, we present POASTA: a fast, memory-efficient, and optimal POA algorithm that computes many fewer alignment states than SPOA, thus enabling the construction of much larger POA graphs (Figure 1). It is built on top of the A* algorithm (Hart et al., 1968), with a new POA-specific heuristic. Inspired by the recently published wavefront algorithm for pairwise alignment (Marco-Sola et al., 2021), it also exploits exact matches between a query sequence and the graph. We additionally introduce a novel superbubble-informed (Onodera et al., 2013) technique for pruning the number of computed alignment states without sacrificing alignment optimality. We benchmarked POASTA against SPOA (Vaser et al., 2017) on diverse sets of bacterial housekeeping genes extracted from RefSeq and demonstrated its increased performance. Additionally, we constructed megabase-length alignments of 342 *Mycobacterium tuberculosis* sequences, demonstrating its reduced memory usage and highlighting POASTA’s ability to align much longer sequences than previously possible.

**Fig. 1.**
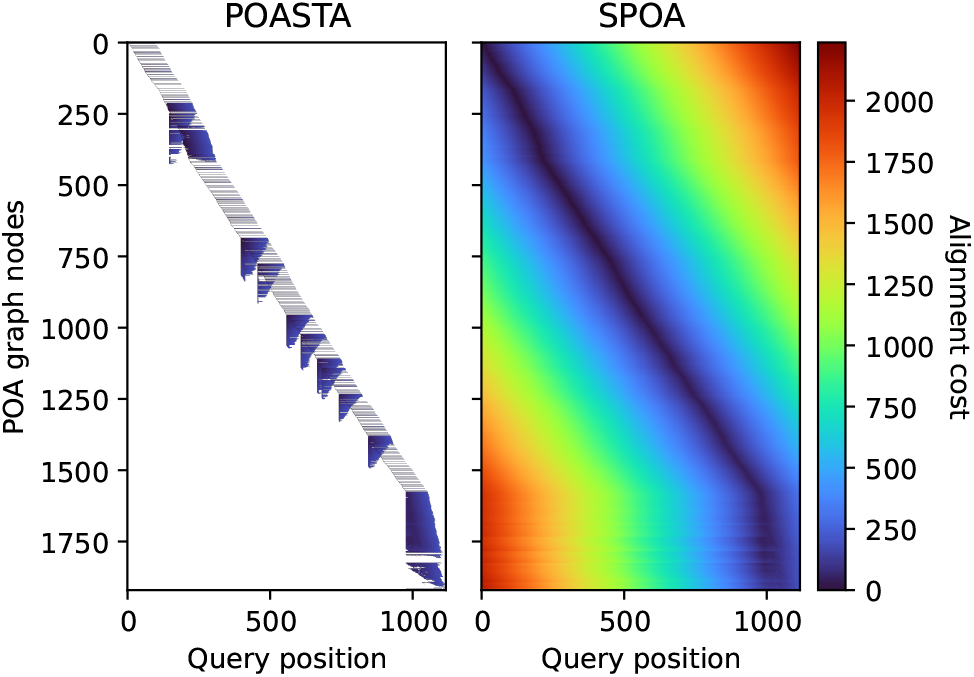
Representation of the dynamic programming matrix to compute the global alignment of a *nusA* gene sequence (x-axis) to a POA graph constructed from 50 other *nusA* gene sequences (y-axis). Each pixel represents a computed alignment state, and the color represents the alignment cost of that state. White pixels represent uncomputed states.

## Methods

POA algorithms compute an MSA by iteratively computing the alignment of a query to a directed acyclic graph (DAG) representing the MSA from the previous iteration (Lee et al., 2002). Instead of the original DP formulation (Supplementary Methods; Supplementary Figure 1a), POASTA’s algorithm is based on an *alignment graph* (Supplementary Figure 1b; not to be confused with the POA graph), enabling the use of common graph traversal algorithms such as the A* algorithm to compute alignments (Hart et al., 1968; Rautiainen and Marschall, 2017; Jain et al., 2020; Ivanov et al., 2020). POASTA further accelerates alignment using three novel techniques: 1) a cheap-to-compute, POA-specific heuristic for the A* algorithm (Figure 2a), 2) a depth-first search component, greedily aligning exact matches between the query and the graph (Figure 2b); and 3) a method to detect and prune alignment states that are not part of the optimal solution, informed by the POA graph topology (Figure 2c). Together they substantially reduce the number of computed alignment states.

**Fig. 2.**
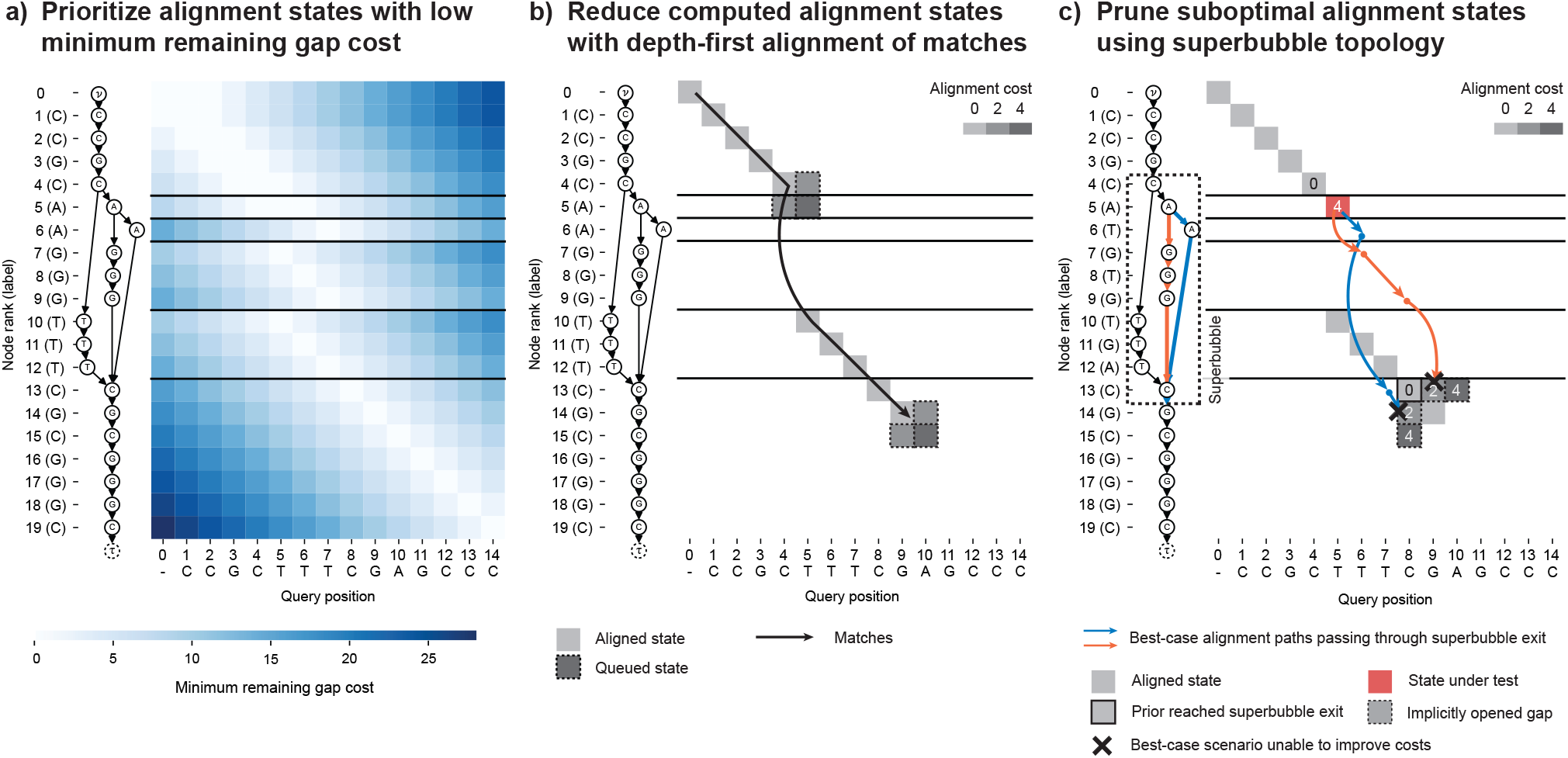
POASTA is based on the A* algorithm and accelerates alignment through three algorithmic innovations: **(a)** A novel heuristic for POA that prioritizes alignment states with a low minimum remaining gap cost (light-colored squares); i.e., states where the unaligned query sequence length is similar to the path lengths to the POA graph end node *τ*. **(b)** Reducing the number of computed alignment states by combining the A* algorithm with a depth-first search component, greedily aligning matches between the query and a path in the graph (black arrow). Adjacent insertion and deletion states are only queued when encountering a mismatch (squares with dashed borders). **(c)** Using knowledge about *superbubble* topology to prune states not part of the optimal solution. POASTA checks whether the best-case alignment paths (blue and orange arrows) from a state under test (red square) can improve over the costs of implicitly opened gaps from prior reached bubble exits (bordered squares).

### Definitions and notation

To describe the algorithm in detail, we will use the following notation. A POA graph *G* = (*V, E*) is a character-labeled DAG, where nodes *v* ∈ *V* represent the symbols in the input sequences, each labeled with a character from an alphabet Σ. Edges (*u, v*) ∈ *E* connect nodes that are adjacent in at least one input sequence. We additionally assume the POA graph has a special start node *ν* with outgoing edges to all nodes with no other incoming edges and a special termination node *τ* with incoming edges from all nodes with no other outgoing edges.

The optimal alignment of a query sequence *Q* = *q*_1_*q*_2_ … *q*_*m*_ (of length *m*) to *G* is the alignment of *Q* to a path *π* = *νv*_1_*v*_2_…*v*_*n*_*τ*, spelling a sequence *R* that minimizes the alignment cost *C* (Supplemental Figure 1a). Commonly used cost models are linear gap penalties and gap-affine penalties. In the former, each gap position is weighted equally, and the alignment cost is defined as *C* = *N*_*m*_Δ_*m*_ + *N*_*x*_Δ_*x*_ + *N*_*g*_Δ_*g*_, where *N*_*m*_ represents the number of matches, *N*_*x*_ is the number of mismatches, and *N*_*g*_ is the total length of gaps. The cost of each alignment operation is represented by Δ_*m*_, Δ_*x*_, and Δ_*g*_, representing the cost of a match, mismatch, and a gap, respectively. In the case of gap-affine penalties, opening a new gap has a different (typically higher) cost than extending an existing gap. The total cost is defined as *C* = *N*_*m*_Δ_*m*_ + *N*_*x*_Δ_*x*_ + *N*_*o*_Δ_*o*_ + *N*_*g*_Δ_*g*_, with *N*_*o*_ the number of distinct gaps and Δ_*o*_ the cost of opening a new gap (Durbin et al., 1998). POASTA supports both the gap-linear and the gap-affine cost models, though it constrains Δ_*m*_ to be zero and all other costs Δ_*x*_, Δ_*o*_, Δ_*g*_ to be *>* 0. In practice, this is not a stringent constraint, as a cost model with non-zero match cost can be reformulated into one with a match cost of zero (Eizenga and Paten, 2022). For clarity, we focus on the gap-linear cost model; the use of POASTA with the gap-affine cost is explained in the Supplemental Methods.

The alignment graph *G*^*A*^ = (*V* ^*A*^, *E*^*A*^) is a product of the POA graph and the query sequence, and paths through it represent possible alignments between them. Nodes ⟨*v, i*⟩ ∈ *V* ^*A*^ = (*V* × {0, 1, …, *m*}) represent *alignment states* with a cursor pointing to a node *v* in the POA graph and a cursor to a query position *i* (Supplementary Figure 1b). Edges in the alignment graph correspond to different alignment operations, such as (mis)match, insertion, or deletion, and are weighted with the respective alignment cost. Edges connect alignment states where either one (indel) or both of the cursors have moved ((mis)match), and the construction of edges is further detailed in the Supplementary Methods. The lowest cost path in the alignment graph from ⟨*ν*, 0⟩ to alignment termination state ⟨*τ, m*⟩ is equivalent to the optimal alignment of *Q* to *G*.

### Optimal alignment with A* using a minimum remaining gap cost heuristic

To compute the lowest-cost path in the alignment graph, i.e., the optimal alignment, POASTA uses the A* algorithm (Hart et al., 1968). For POASTA, we adapted the widely used gap-cost heuristic for pairwise alignment to POA (Figure 2a) (Ukkonen, 1985; Hadlock, 1988). This heuristic is *admissible*, i.e., it is a lower bound on the true remaining cost, thus guaranteeing that A* finds the lowest-cost path. The intuition behind the heuristic is to prioritize alignment states in which the length of the unaligned query sequence is similar to the path lengths to the end node *τ*.

To compute heuristic *h*⟨*v, i*⟩, POASTA scans the POA graph before alignment starts and stores the shortest and longest path length to the end node *τ* for all nodes in the graph, denoted as 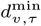 and 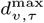. This can be computed in *O*(*V* + *E*) time by visiting the nodes in reverse topological order. POASTA compares these path lengths to the length of the unaligned query sequence *l*_*r*_ = *m* − *i* and infers the minimum number of indel edges to traverse from ⟨*v, i*⟩ to the alignment termination ⟨*τ, m*⟩ state as follows:

#### Definition 1

(Minimum number of indel edges)

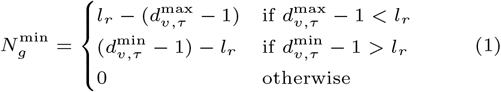

We subtract one from 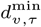 and 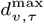 to exclude the edge towards *τ*.

*Proof* See Supplemental Methods. □

Combining the computed minimum number of indel edges to traverse with the alignment cost model, e.g., the linear gap cost model, enables us to compute the heuristic.

#### Definition 2

(Minimum remaining gap cost heuristic)

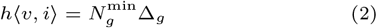

#### Lemma 1

(Admissibility) *h*⟨*v, i*⟩ *is admissible*.

*Proof* The true remaining alignment cost, using linear gap penalties and assuming a match cost Δ_*m*_ of zero, is defined as *C*_*r*_ = *N*_*x*_Δ_*x*_ + *N*_*g*_Δ_*g*_, where *N*_*x*_ and *N*_*g*_ represent the number of remaining mismatches and the total remaining gap length respectively, Δ_*x*_ the mismatch cost, and Δ_*g*_ the gap cost.

Using Definition 1, we infer that 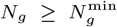. Since the mismatch cost Δ_*x*_ ≥ 0, we note that the *N*_*x*_Δ_*x*_ ≥ 0, and thus observe that

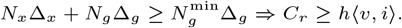

*h*⟨*v, i*⟩ is thus a lower bound on the true remaining alignment cost. □

### Depth-first alignment of exact matches between query and graph

To further speed up alignment and reduce the number of computed alignment states, POASTA greedily aligns exact matches between the query and graph (Figure 2b). This is possible because POASTA requires that the alignment cost for a match is zero, and all other alignment costs are ≥ 0. Traversing a match edge ⟨*u, i*⟩ → ⟨*v, i* + 1⟩ will always be the optimal choice if the latter state has not been visited yet since match edges have zero cost and all other paths (requiring indels) will have higher or equal cost (Ivanov et al., 2020; Marco-Sola et al., 2021). This implies that in the presence of an unvisited match, we can ignore insertion edge ⟨*u, i*⟩ → ⟨*u, i* + 1⟩ and deletion edge ⟨*u, i*⟩ → ⟨*v, i*⟩.

To implement this, POASTA combines the regular A* algorithm with a depth-first search (DFS) component. When a state ⟨*u, i*⟩ is popped from the A* queue, we initiate a DFS from this state. We assess whether a successor state ⟨*v, i* + 1⟩ ∀*v* : (*u, v*) ∈ *E* is a match; if it is, we push it on the stack to be processed in the next DFS iteration; when there is a mismatch, we append it to the A* queue. In the latter case, we no longer can ignore the insertion and deletion edges, so we additionally queue insertion state ⟨*u, i* + 1⟩, and deletion state ⟨*v, i*⟩. Using DFS enables greedily aligning long stretches of exact matches, even in the presence of branches in the graph.

### Pruning alignment states not part of the optimal solution

When POASTA’s depth-first alignment finds a long stretch of matching sequence, the corresponding path through the POA graph might traverse a *superbubble* (Onodera et al., 2013). A superbubble (*s, t*) is a substructure in the POA graph with specific topological features (Supplementary Figure 2): it is acyclic; it has a single entrance *s* and a single exit *t*; all paths leaving *s* should end in *t*; and no path from “outside” the superbubble can have an endpoint inside the bubble. The set of nodes *U* on paths from *s* to *t* is called the *interior* of a bubble, which can be empty. In a POA graph, superbubbles represent the different alleles present at particular loci in the MSA.

POASTA exploits the fact that all paths through a superbubble have a common endpoint, its exit *t*. If an alignment state ⟨*t, p*⟩ is reached during alignment with a particular cost *C*^*t,p*^, POASTA can detect whether another yet-to-visit state ⟨*v, i*⟩ : *v* ∈ *U* ∪ {*s*} that is part of the same superbubble, can improve over this cost. This is especially effective when combined with the depth-first greedy alignment described above; if a bubble exit is reached at a low cost because of a long stretch of matching sequence, we can often prune alignment states on alternative paths through the bubble because they can not improve over the already-found path.

To quickly retrieve topological information about super-bubbles, POASTA constructs a *superbubble index* before alignment. For every node in the POA graph, it stores the superbubbles in which it is contained, along with the shortest and longest path length to the corresponding superbubble exit (Figure 3a). For example, the red node (node 5) in the example shown in Figure 3b has two paths to the superbubble exit (node 13): one path with length 2 (blue) and one path with length 4 (orange). POASTA identifies superbubbles using the *O*(*V* + *E*) algorithm described by Gärtner et al. (2018). The shortest path lengths can be computed using a backward breadth-first search (BFS), and the longest path lengths can be computed by recursively visiting nodes in postorder, both *O*(*V* + *E*) operations.

**Fig. 3.**
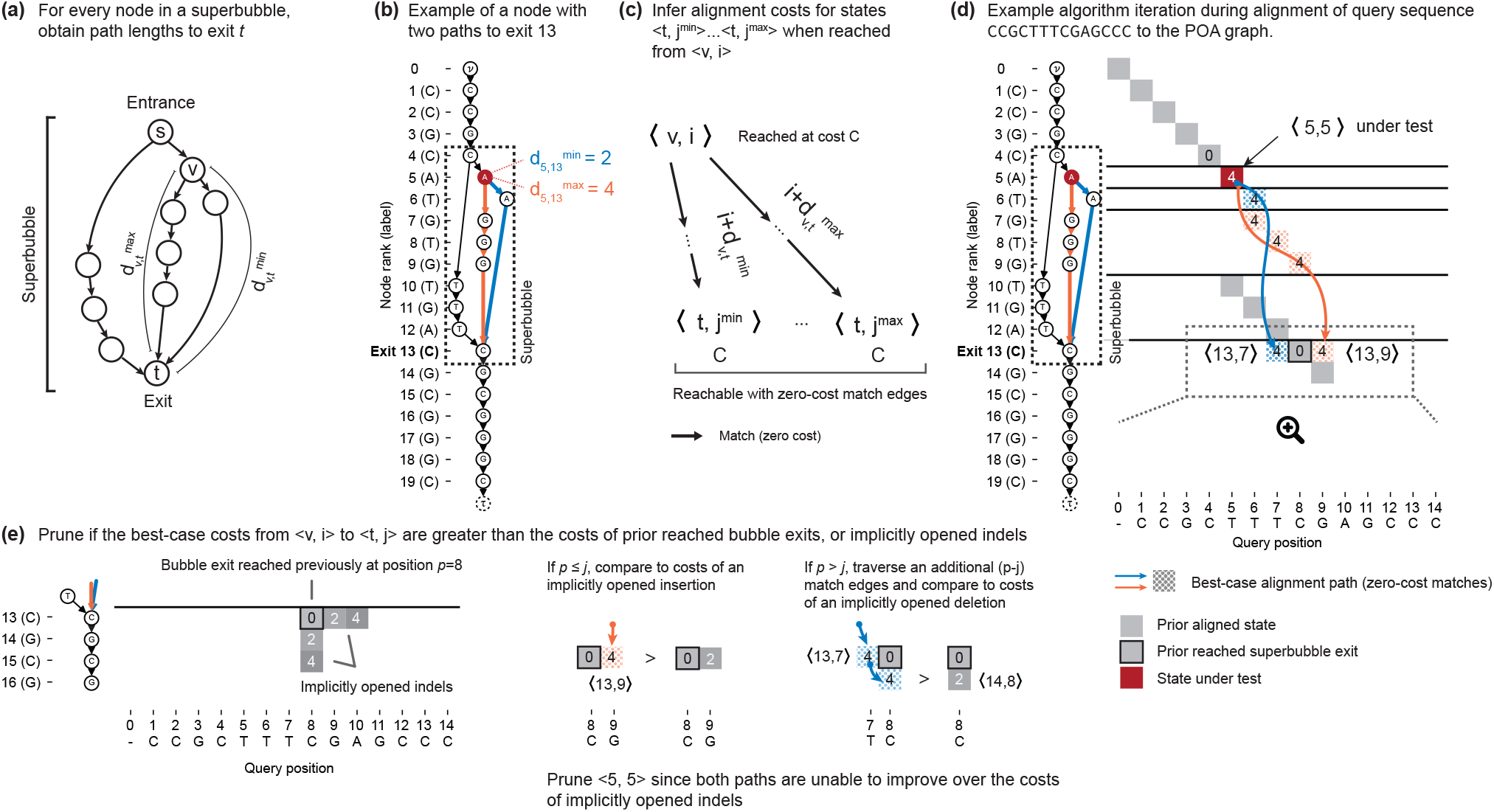
POASTA detects and prunes alignment states that are not part of the optimal solution. **(a)** A superbubble with entrance *s* and exit *t*. For every node *v* in the superbubble, POASTA stores the minimum and maximum path length, 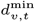, and 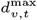, to exit *t*. **(b)** An example POA graph with a superbubble (dashed rectangle) and the path lengths from the highlighted node 5 (red) to the superbubble exit (orange and blue paths). **(c)** The path lengths to the superbubble exit are used during alignment to infer the range of states reachable with zero-cost match edges (black arrows) from another state ⟨*v, i*⟩. **(d)** An example aligning the query CCGCTTTCGAGCCC to the graph in (b). Grey squares: states aligned in a prior iteration. Red square: state under test. Blue and orange arrows and dotted squares: best-case alignment paths from the state under test to the superbubble exit. **(e)** POASTA compares best-case alignment costs from ⟨*v, i*⟩ (blue and orange dotted squares) to implicitly opened indels from prior reached bubble exits (grey squares). Implicitly opened indels act as an upper bound for the alignment cost of yet-to-visit states.

To test if a state ⟨*v, i*⟩ should be pruned, POASTA first uses the superbubble index to infer the range of states ⟨*t, j*^min^⟩ … ⟨*t, j*^max^⟩ reachable from ⟨*v, i*⟩ assuming the best-case scenario of traversing zero-cost match edges (Figure 3c). For example, when aligning a query CCGCTTTCGAGCCC to the graph in Figure 3b, POASTA will initially find a long stretch of matches between the query and a path in the graph, traversing the superbubble (4, 13) (Figure 3d; grey squares). In a following iteration, it tests alignment state ⟨5, 5⟩, where node 5 is part of the same superbubble (4, 13), which is reached with an alignment cost 4 (Figure 3d; red square). It looks up the path lengths to the superbubble exit 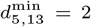 and 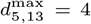 and infers that we can reach ⟨13, 7⟩ and ⟨13, 9⟩ from ⟨5, 5⟩ with the same alignment cost of four (Figure 3d; blue and orange arrows and dotted squares).

POASTA can now compare this best-case alignment cost, when reached from a state ⟨*v, i*⟩, to the alignment costs of states that reached the superbubble exit prior, or an implicitly opened gap from those. Implicitly computed gap costs will act as an upper bound on the costs for yet-to-visit alignment states (Figure 3e). For example, the orange path in Figure 3d could reach alignment state ⟨13, 9⟩ with an alignment cost of four. However, alignment state ⟨13, 9⟩ is also reachable from the prior reached bubble exit ⟨13, 8⟩, by opening an insertion and reaching it with a lower cost of two (Figure 3e). Similarly, the blue path in Figure 3d could reach alignment state ⟨13, 7⟩ with an alignment cost of four. This state has not yet been reached and is also not reachable by opening a gap from a previously reached exit. However, if we extend the blue path, assuming additional traversal of zero-cost match edges, we reach an alignment state ⟨14, 8⟩ which is reachable from a previously reached exit by opening a deletion. The opened deletion would reach ⟨14, 8⟩ with a cost of two, lower than the cost of four when reached through the blue path. Since both best-case scenarios from ⟨5, 5⟩ would result in higher alignment costs compared to opened indels from a prior reached exit, POASTA infers ⟨5, 5⟩ will not be part of the optimal solution and prunes it from further consideration.

In the example discussed above, the bubble exit was only reached once. Bubble exits, however, can be reached multiple times during alignment (with varying alignment costs). When testing whether a state can be pruned, all previously reached positions should be considered (Supplementary Figure 3). The Supplemental Methods describe the details of how POASTA prunes alignment states when the bubble exit has been reached multiple times.

## Results

### Benchmarking using bacterial housekeeping genes

To compare POASTA’s speed and memory usage to the current state of the art, we generated multiple benchmark datasets from bacterial housekeeping genes (*dnaG, nusA, pgk, pyrG*, and *rpoB*). These genes are present in nearly all bacteria and are commonly used to create bacterial phylogenies, requiring MSA (Wu and Eisen, 2008). We downloaded all 40,188 RefSeq-complete genomes representing the breadth of bacterial diversity and extracted genes of interest using the accompanying gene annotations. Gene sequences were deduplicated and coarsely clustered using single-linkage hierarchical clustering. This resulted in multiple genus-spanning clusters. For each gene family, we selected one or more clusters as benchmark datasets, choosing clusters with at least 100 sequences and varying pairwise ANI (Supplemental Methods). The 13 selected benchmark sets each contained 140-2,385 gene sequences, with mean sequence lengths of 1-4kbp and pairwise ANIs of 82%-97% (Supplemental Table 1).

### POASTA constructs multiple sequence alignments 4x faster than other methods

We assessed POASTA’s runtime and memory compared to SPOA. We only compared against SPOA since it is the only other POA algorithm that guarantees optimal partial order alignments. We ran both tools to compute the full MSA of the 13 selected datasets and recorded their total runtime and memory usage.

For 12 of 13 datasets, POASTA computed the complete MSA faster than SPOA, achieving an average speed-up of 4.1x. The highest speed-up was 9.8x (Figure 4a). The one instance where SPOA was faster corresponded to the gene set with the lowest pairwise ANI (82.6%). POASTA’s strongest relative performance was in settings with ANIs of 90-100% and sequences longer than 1,500 bp (Figure 4b,c). Furthermore, POASTA used less memory than SPOA, requiring, on average, 2.6x less memory than SPOA (Figure 4d-f).

**Fig. 4.**
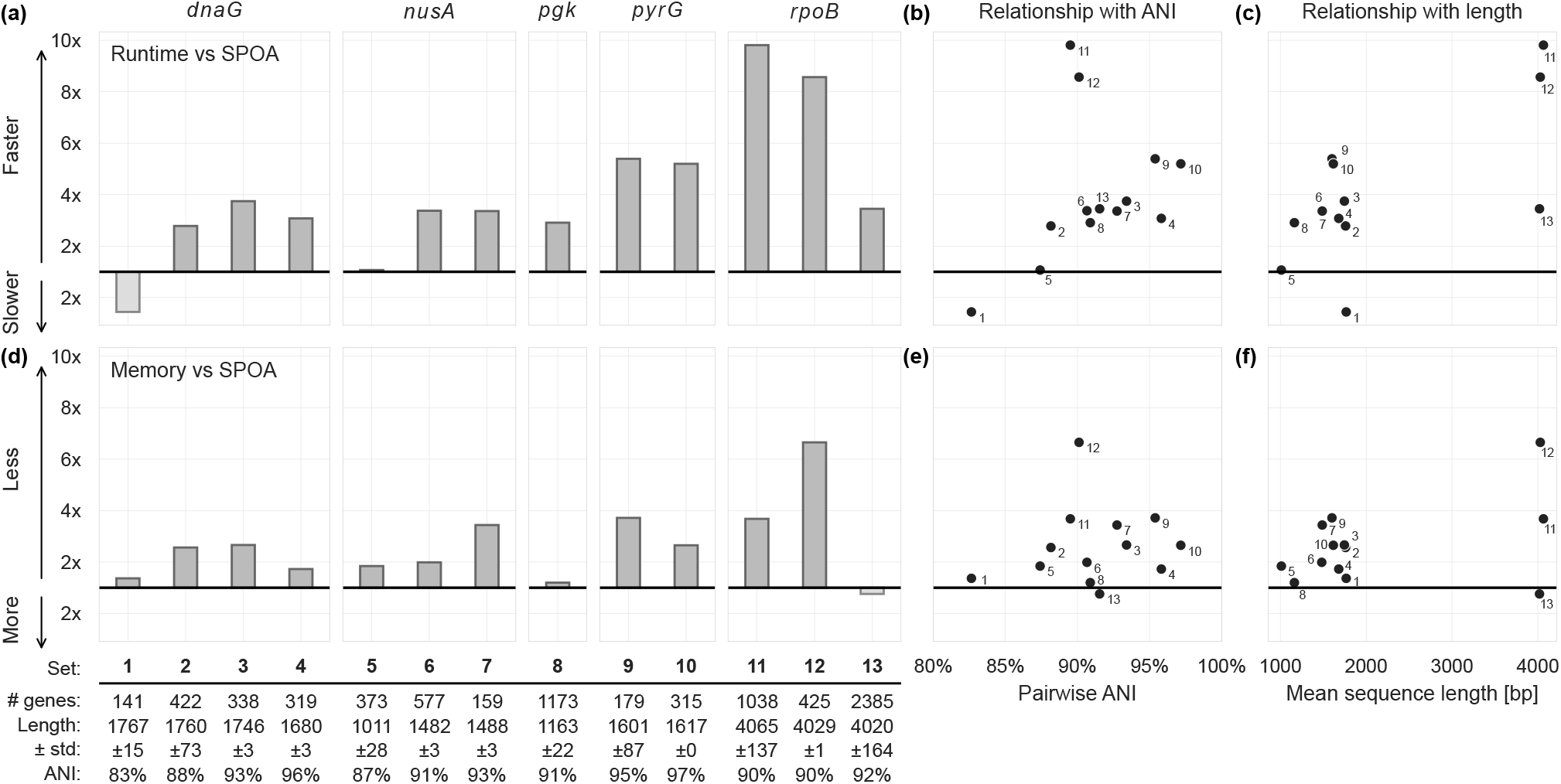
POASTA creates multiple sequence alignments of sequences from five bacterial housekeeping genes, in all but one case faster than SPOA and with less memory. **(a)** Relative runtime of POASTA compared to SPOA for each set of gene sequences. **(b)** The relationship between pairwise ANI of each gene sequence set and POASTA’s relative runtime. **(c)** The relationship between mean sequence length and POASTA’s relative runtime. **(d)** Relative memory usage of POASTA compared to SPOA for each set of gene sequences. **(e)** The relationship between pairwise ANI of each sequence set and POASTA’s relative memory usage. **(f)** The relationship between the mean sequence length of each sequence set and POASTA’s relative memory usage.

### POASTA enables the construction of megabase-length POA graphs

To further test POASTA’s limits, we benchmarked its ability to align datasets with average sequence lengths of approximately 250 kbp, 500 kbp, and 1 Mbp. We extracted subsequences from all 370 RefSeq-complete whole genome assemblies of *Mycobacterium tuberculosis*, covering a broad range of the species’ diversity (including representatives from all known lineages; Mash-estimated average pairwise ANI of 99.3% (Ondov et al., 2016)). *M. tuberculosis* has relatively little large-scale structural variation, including few large inversions or genes translocating to different locations, which POA cannot model and align accurately. After orienting genomes such that each started with the gene *dnaA*, we truncated at specific shared genes to achieve sequences of the desired length (Supplemental Methods). For the 250 kbp, 500 kbp, and 1 Mbp benchmarks, we truncated at the genes *trmB, thiE*, and *gltA2*, respectively. Since POA expects sequences to be colinear, we also excluded 28 genomes with more than 15% (≥ 660 kbp) of its complete genome inverted with respect to the canonical reference H37Rv (Supplemental Methods).

POASTA successfully computed MSAs for the 250 kbp, 500 kbp, and 1 Mbp benchmark sets with manageable runtimes and memory (Table 1). None of these alignments could be completed with SPOA, which required more memory than the 240 GB available in the Google Cloud VM used for benchmarking (Supplemental Methods). The estimated memory requirements for SPOA would be 0.95, 3.5, and 13 TB for the 250, 500, and 1,000 kbp benchmarks, respectively (assuming 32-bit integers for storing scores).

**Table 1.** POASTA runtime and peak memory usage for three benchmark sets comprising 342 *M. tuberculosis* sequences of approximately 250, 500, and 1,000 kbp.

| Sequence set | Runtime | Max. memory |
| --- | --- | --- |
| 250 kbp | 5.3 h | 63.8 GB |
| 500 kbp | 24 h | 120 GB |
| 1,000 kbp | 69 h | 231 GB |

We assessed computed alignments at a known drug resistance locus to validate that the MSA correctly captured known variation. In *M. tuberculosis*, the S450L change in the *rpoB* gene is one of the most common rifampicin resistance-causing mutations (Munir et al., 2019; Jamieson et al., 2020). We first characterized codons representing the 450th amino acid of *rpoB* using just the reference genomes and accompanying gene annotations. We obtained each codon using the start position of the *rpoB* gene to compute the reference locus representing the 450th amino acid of *rpoB*. In our set of genomes, we similarly observed that the S450L mutation is the most common allele present other than the reference or wild-type allele (Table 2; 103 genomes have the S450L mutation). To check if the observed codons were correctly aligned in the POA graph, we extracted a small subgraph surrounding the 450th amino acid of *rpoB* in H37Rv (Figure 5). While this subgraph was obtained using H37Rv coordinates, all codons listed in Table 2 were also represented as different paths in the graph, and the graph edge counts, indicating the number of genomes sharing that edge, matched the codon counts obtained through gene annotations. POASTA thus correctly captured known variation at this locus while the alignments were computed unaware of genes.

**Table 2.** Diversity of codons across 342 *M. tuberculosis* genomes representing the 450th amino acid in the *rpoB* gene. In three genomes, there was uncertainty about the second base in the triplet indicated by IUPAC code N (any base) or Y (C or T).

| Codon | Amino acid | Count |
| --- | --- | --- |
| TCG (reference) | S | 232 |
| TTG | L | 103 |
| TTT | F | 2 |
| TNG | - | 2 |
| GCG | A | 1 |
| TGG | W | 1 |
| TYG | - | 1 |

**Fig. 5.**
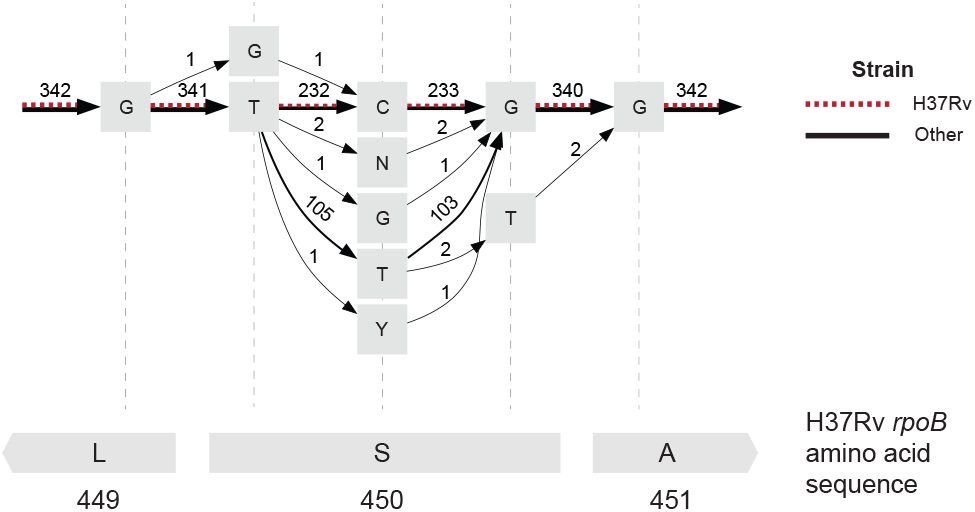
The POA subgraph surrounding the 450th amino acid in *M. tuberculosis* H37Rv *rpoB* (red dashed edges) captures extensive allelic diversity in other references (black edges). Grey squares represent nodes in the POA graph labeled with a base or an IUPAC code representing uncertainty about the base at that site (N: any base, Y: C or T). Edges are labeled with the number of genomes that share that edge. The bottom grey rectangles represent the H37Rv amino acid sequence.

## Discussion

In this work, we introduced POASTA, an optimal POA algorithm supporting gap-affine penalties with increased performance. These improvements are achieved using three algorithmic innovations: a minimum remaining gap cost heuristic for A*, depth-first greedy alignment of matches, and pruning states not part of the optimal solution using superbubble topology. In benchmarking on short sequences (1-4 kbp), POASTA was, on average, 4.1x faster than the current state-of-the-art SPOA (Vaser et al., 2017) and used 2.6x less memory. On longer sequences (250-1,000 kbp), POASTA generated alignments with manageable runtime and memory, while SPOA failed.

POASTA includes several algorithmic innovations inspired by recent advances in pairwise and graph alignment. For example, POASTA takes inspiration from the recently published wavefront algorithm (WFA), a fast algorithm for pairwise alignment (Marco-Sola et al., 2021). WFA similarly exploits exact matches between sequences and rapidly computes alignments by only considering the furthest-reaching points on DP matrix diagonals. However, their DP matrix diagonal formulation does not directly apply to graph alignment. In contrast to pairwise alignment, a stretch of exact matches between the query and the graph may span multiple diagonals in the DP matrix because of branches in the graph, complicating the definition of furthest-reaching points. While others have introduced variants of the WFA for graphs (Zhang et al., 2022; Holt et al., 2023), none support the gap-affine scoring model, which is preferred because it gives more biologically relevant alignments Durbin et al. (1998). As an alternative to processing only the furthest-reaching points on a diagonal, POASTA uses its knowledge of graph topology, as stored in its superbubble index, to detect and prune alignment states that are not part of the optimal solution, thus speeding up alignment.

POASTA additionally takes inspiration from the recent read-to-graph aligner Astarix. Like POASTA, Astarix uses the A* algorithm for alignment, though with a different heuristic (Ivanov et al., 2020, 2022). The benefit of our minimum remaining gap cost A* heuristic is the simplicity of the required computation. All preprocessing can be done in *O*(*V* + *E*) time, and all the necessary data is stored in *O*(*V*) additional memory. The fast computation of the heuristic is important because the POA graph is updated at each iteration. Combined, these innovations can substantially reduce the number of computed alignment states, speeding up the construction of the complete MSA and enabling MSAs for longer sequences than was previously possible.

POASTA did not improve over SPOA in every scenario—it performed less well than SPOA in settings with high sequence diversity, where there are fewer stretches of exact matches for POASTA to exploit. In this situation, POASTA must explore more mismatch and indel states, increasing computation time. Though POASTA still computes fewer alignment states than SPOA, its runtime can become longer because the A* algorithm is less predictable and less CPU cache-efficient compared to computing the full DP matrix row-by-row in a contiguous block of memory. Despite POASTA’s higher compute time *per alignment state* compared to SPOA, the reduction in computed alignment states is often large enough to still gain a net speed increase.

We envision several future improvements to the POASTA algorithm. Bi-directed variants of the A* algorithm, where the search for the shortest path is started from both the start and the end, could substantially improve POASTA’s runtime with respect to sequence diversity (de Champeaux, 1983). A more informative A* heuristic, e.g., the recently published seed-heuristic (Ivanov et al., 2022), could speed up alignment by improving estimates of the remaining alignment cost, resulting in improved prioritization of alignment states to visit. Other strategies could be to utilize GPUs since massively parallel versions of A* exist (Zhou and Zeng, 2015). Finally, we could combine the superbubble index with the Gwfa algorithm (Zhang et al., 2022) to link diagonals across nodes and increase power to prune suboptimal alignment states.

## Conclusions

We present POASTA, a novel optimal algorithm for POA. Through several algorithmic innovations, POASTA computed the complete MSA faster than existing tools in diverse bacterial gene sequence sets. It further enabled the creation of much longer MSAs, as demonstrated by successfully constructing MSAs from *M. tuberculosis* sequence sets with average sequence lengths of up to 1 Mbp. The algorithms and ideas presented here will accelerate the development of scalable pangenome construction and analysis tools that will drive the coming era of genome analysis.

## Acknowledgments

We would like to thank Fabio Cunial and Ryan Lorig-Roach for their helpful discussions and their reviews of early versions of the manuscript. This project has been funded in part with federal funds from the National Institute of Allergy and Infectious Diseases, National Institutes of Health, Department of Health and Human Services, under Grant Number U19AI110818 to the Broad Institute.

## Availability of data and materials

POASTA is written in Rust and available under the BSD-3-clause license at https://github.com/broadinstitute/poasta (DOI: 10.5281/zenodo.11153323). POASTA is available as both a standalone utility and a Rust crate that can be included as part of other software packages. The benchmark suite is written in Rust and Python and is available under the same license at https://github.com/broadinstitute/poa-bench. The data underlying this paper are included in the benchmark suite repository (DOI: 10.5281/zenodo.11153368).

