## Supplemental Figures and Text for "Fast and exact gap-affine partial order alignment with POASTA"

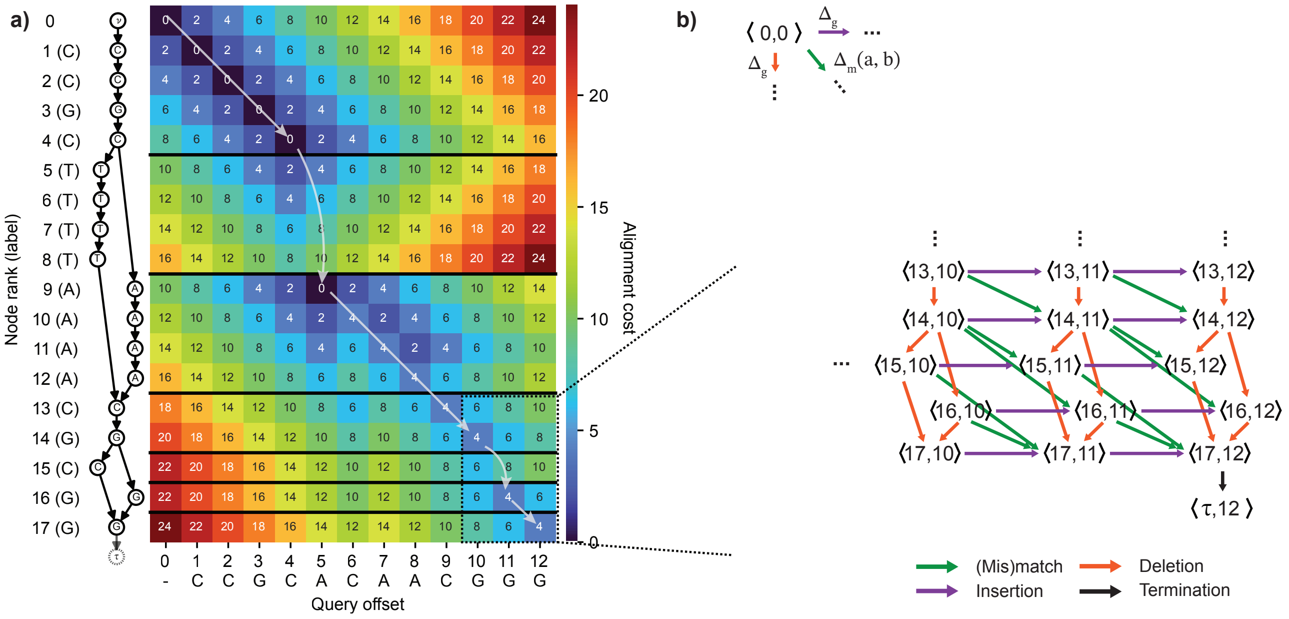


**Fig. 1. (a)** Example computation of aligning "CCGCACAACGGG" to a POA graph, with mismatch cost $\Delta_x = 4$, and gap cost $\Delta_g = 2$. The white arrows indicate the optimal alignment path. **(b)** A subgraph of the full *alignment graph*, corresponding to POA graph nodes 13-17, and query offset 10-12. A node $\langle v, o\rangle$ in the alignment graph represents a cursor to a node in the POA graph $v$ and a query offset $o$. The various alignment operations ((mis)match, insertion, deletion) correspond to different kinds of edges. Insertion and deletion edges are weighted with the gap cost $\Delta_g$, and (mis)match edges with a function $\Delta_m(a, b) = \{\Delta_x$ if $a \neq b$, and 0 otherwise.


© The Author 2024. Published by Oxford University Press. All rights reserved. For permissions, please

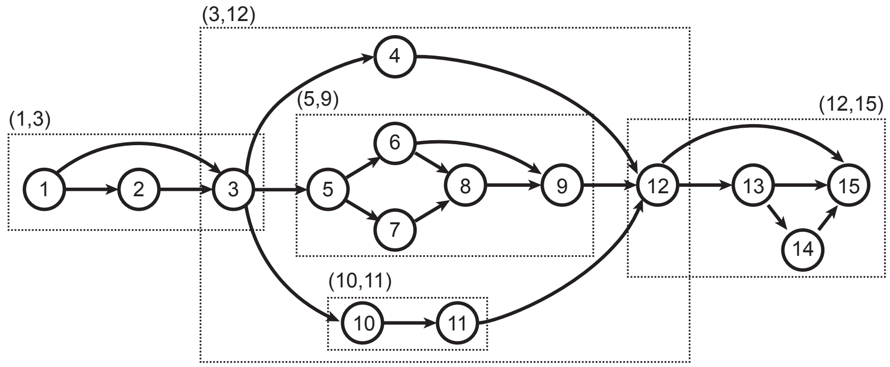


**Fig. 2.** An example graph containing multiple superbubbles. Each superbubble is marked with a dotted rectangle, labeled with its (*entrance*, *exit*). Superbubbles can be nested within each other: superbubbles (5, 9) and (10, 11) are contained within superbubble (3, 12). Superbubble (10, 11) is an example of a superbubble without an interior.

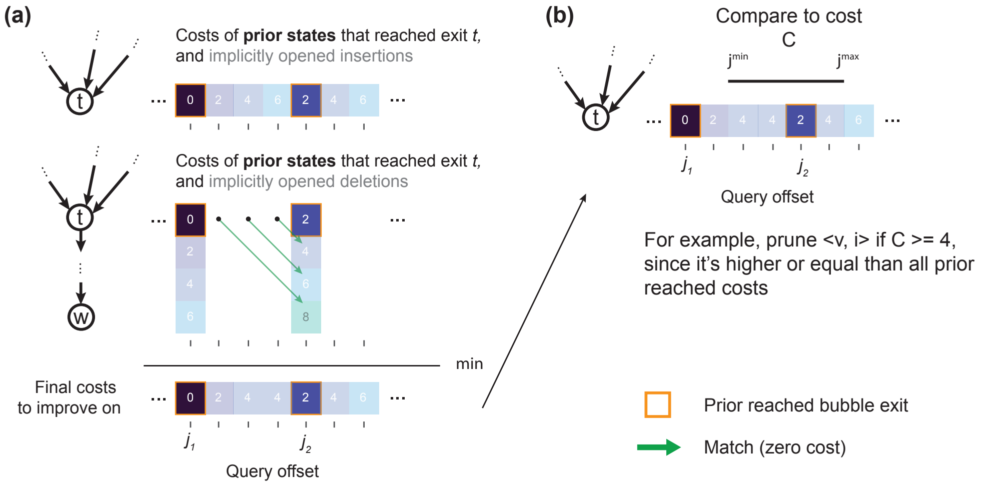


**Fig. 3.** POASTA considers all prior reached bubble exits when testing if a state can be pruned. **(a)** Example that shows alignment states that reached exit $t$ previously at query position $j_1$ and $j_2$ (dark, orange-bordered squares). We also display implicitly opened insertions (top; light-colored squares) and implicitly opened deletions (middle). The costs of a deletion, reaching some node $w$ downstream of $t$, can be linked to an alignment state involving $t$ by tracing back the assumed best-case path of matches (green arrows). The final costs another state needs to improve upon are obtained by taking the minimum (bottom). **(b)** If none of the alignment paths from a state $\langle v, i \rangle$ reaching $\langle t, j^{\min} \rangle \ldots \langle t, j^{\max} \rangle$ with a cost $C$ can improve over implicitly opened indels, that state will be pruned.

### Supplemental Methods

#### The connection between dynamic programming recurrence and the alignment graph

##### *Gap-linear alignment costs*

The conventional dynamic programming (DP) recurrence for POA with gap-linear costs is (Lee et al., 2002):

$$S_{v,i} = \min \begin{cases} S_{u,i-1} + \Delta(\sigma(v), q_i) & \forall u : (u,v) \in E \\ S_{u,i} + \Delta_g & \forall u : (u,v) \in E \\ S_{v,i-1} + \Delta_g \end{cases} \tag{1}$$

Here, $\sigma(v) \rightarrow \Sigma$ returns the node label for a node $v$, and $\Delta(a,b) \rightarrow \mathbb{Z}$ is a function that returns the match cost $\Delta_m$ if $a = b$, and mismatch cost $\Delta_x$ if $a \neq b$. The three cases correspond to a (mis)match between the graph and query, opening or extending a deletion, and opening or extending an insertion.

To translate the recurrence to edges in an alignment graph, we define the edge set $E^A$ as follows. Edges connect two alignment states $\langle u, i\rangle \rightarrow \langle v, j\rangle$ if one of the following conditions hold:

- **Match and mismatch.** $(u,v) \in E$, $v \neq \tau$, and $i+1 = j$, $j \leq m$, with the (mis)match cost $\Delta(\sigma(v), q_j)$ as weight;
- **Deletion.** $(u,v) \in E$, $v \neq \tau$, and $i = j$, with the gap cost $\Delta_g$ as weight;
- **Insertion.** $u = v$, $u, v \neq \tau$, and $i+1 = j$, $j \leq m$, with the gap cost $\Delta_g$ as weight;
- **Termination.** $(u,v) \in E$, $v = \tau$, and $i = j = m$, with zero cost.

These edges (except for the termination edge) are analogous to the different cases in Equation 1.

##### *Gap-affine alignment costs*

To compute the gap-affine alignment, the Smith-Waterman-Gotoh (SWG) DP recurrence for affine pairwise alignment (Gotoh, 1982) can be adapted to POA as follows:

$$\begin{cases} I_{v,i} & = \min\{M_{v,i-1} + \Delta_o + \Delta_g,\ I_{v,i-1} + \Delta_g\} \\ D_{v,i} & = \min\{M_{u,i} + \Delta_o + \Delta_g,\ D_{u,i} + \Delta_g\} \\ & \qquad \forall u : (u,v) \in E \\ M_{v,i} & = \min\{I_{v,i},\ D_{v,i},\ M_{u,i-1} + \Delta(\sigma(v), q_i)\} \\ & \qquad \forall u : (u,v) \in E \end{cases} \tag{2}$$

The two cases for $I_{v,i}$ correspond to opening an insertion and extending an insertion; the two cases for $D_{v,i}$ correspond to opening a deletion and extending a deletion; and the three cases for $M_{v,i}$ correspond to closing an insertion, deletion, or a (mis)match.

To extend the alignment graph formulation to the gap-affine model, with gap open cost $\Delta_o$ and gap extend cost $\Delta_g$, we define the node set of the gap-affine alignment graph as follows: $V^A = (V \times \{0, \ldots, m\} \times \{M, D, I\})$. In other words, for each pair $v \in V, i \in [0, m]$, we now have three possible alignment states: $\langle v, i, M\rangle$, $\langle v, i, D\rangle$, $\langle v, i, I\rangle$, representing the match, deletion, and insertion state, respectively. Edges in the gap-affine alignment graph are defined as follows:

- **Edges ending in the insertion state**
  - $\langle u, i, M\rangle \rightarrow \langle u, i+1, I\rangle$, $u \neq \tau$, $i+1 \leq m$, weighted with gap open cost $\Delta_o + \Delta_g$
  - $\langle u, i, I\rangle \rightarrow \langle u, i+1, I\rangle$, $u \neq \tau$, $i+1 \leq m$, weighted with gap extend cost $\Delta_g$
- **Edges ending in the deletion state**
  - $\langle u, i, M\rangle \rightarrow \langle v, i, D\rangle$, $(u,v) \in E$, $v \neq \tau$, weighted with gap open cost $\Delta_o + \Delta_g$
  - $\langle u, i, D\rangle \rightarrow \langle v, i, D\rangle$, $(u,v) \in E$, $v \neq \tau$, weighted with gap extend cost $\Delta_g$
- **Edges ending in the (mis)match state**
  - $\langle u, i, M\rangle \rightarrow \langle v, i+1, M\rangle$, $(u,v) \in E$, $v \neq \tau$, $i+1 \leq m$, with (mis)match cost $\Delta(\sigma(v), q_{i+1})$
  - $\langle u, i, I\rangle \rightarrow \langle u, i, M\rangle$, $u \neq \tau$, weighted with zero cost
  - $\langle u, i, D\rangle \rightarrow \langle u, i, M\rangle$, $u \neq \tau$, weighted with zero cost
- **Termination edges**
  - $\langle u, m, M\rangle \rightarrow \langle \tau, m, M\rangle$, $(u, \tau) \in E$, weighted with zero cost

These edges are analogous to the cases in Equation 2.

#### Proof of minimum number of indel edges

Given an alignment state $\langle u, i\rangle$, let $d^{\min}_{u,\tau}$ and $d^{\max}_{u,\tau}$ be the minimum and maximum path length in the POA graph from $u$ to end node $\tau$. We additionally compute the length of the unaligned query sequence $l_r = m - i$. The minimum number of indel edges to traverse is then:

**Definition 1** (Minimum number of indel edges)

$$N^{\min}_g = \begin{cases} l_r - (d^{\max}_{u,\tau} - 1) & \text{if } d^{\max}_{u,\tau} - 1 < l_r \\ (d^{\min}_{u,\tau} - 1) - l_r & \text{if } d^{\min}_{u,\tau} - 1 > l_r \\ 0 & \text{otherwise} \end{cases} \tag{3}$$

We subtract one from $d^{\min}_{u,\tau}$ and $d^{\max}_{u,\tau}$ to exclude the edge towards $\tau$.

*Proof* Let $\mathcal{W} \subset V$ be the subset of POA graph nodes with an outgoing edge to $\tau$, i.e., $\mathcal{W} = \{w : (w, \tau) \in E\}$. By definition of the alignment graph, the alignment termination state is only reachable from alignment states $\langle w, m\rangle : w \in \mathcal{U}$. We will prove each individual case separately.

In the first case, $d^{\max}_{u,\tau} - 1 < l_r$. By definition of $\mathcal{W}$, $\exists w \in \mathcal{W}$ such that $d_{u,w} = d^{\max}_{u,\tau} - 1$, i.e., excluding the last edge towards $\tau$ from the maximum path length. The presence of this maximum length path in the POA graph implies a corresponding path of all (mis)match edges in the alignment graph, reaching the alignment state $\langle w, j\rangle, w \in \mathcal{W}, j = i + d_{v,w}$. Since this traversed the maximum length path in the POA graph, $j$ is also the maximum query position reachable from $\langle u, i\rangle$. Since $d_{u,w} < l_r$, we infer that $j < m$. This means that the alignment termination state is not reachable from $\langle u, j\rangle$, and at least $m - j = l_r - (d^{\max}_{v,\tau} - 1)$ insertion edges need to be traversed to be able to reach the alignment termination state.

In the second case, $d^{\min}_{v,\tau} - 1 > l_r$. Similarly as above, $\exists w \in \mathcal{W}$, such that $d_{u,w} = d^{\min}_{u,\tau} - 1$, i.e., excluding the last edge towards $\tau$ from the minimum path length. To reach the alignment termination state from $\langle u, i\rangle$, we need to traverse at least $d_{u,w}$ (mis)match or deletion edges, since this is the minimum length path to the POA end node. We can, however, traverse only $l_r$ (mis)match edges, since no (mis)match edges exist that would move the query position beyond the query sequence length $m$. After traversing $l_r$ (mis)match edges, we would reach some state $\langle v, m\rangle$, with $v$ being a node on the minimum path in the POA graph $u \to \ldots \to v \to \ldots \to w \to \tau$. To be able to reach the alignment termination state, we need to traverse at least $d^{\min}_{v,\tau} - 1 - l_r$ deletion edges.

In the last case, $d^{\min}_{v,\tau} - 1 < l_r < d^{\max}_{v,\tau} - 1$, which implies that there exist a path from $\langle u, i\rangle$ to $\langle \tau, m\rangle$ without the need to traverse any indel edges. □

#### Extension of the minimum gap cost heuristic function to the gap-affine model

To compute the minimum gap cost heuristic using gap-affine model, we need to take into account that insertion or deletion states do not need to incur the gap-open cost again.

A state $\langle v, i, M\rangle$ always needs to incur the gap-open cost, thus the heuristic is computed as follows:

**Definition 2** $$h\langle v, i, M\rangle = \begin{cases} 0 & \text{if } N^{\min}_g = 0 \\ \Delta_o + N^{\min}_g \Delta_g & \text{otherwise} \end{cases}$$

A state $\langle v, i, I\rangle$ is already in insertion state and would not have to incur the gap open cost again if $d^{\max}_{v,\tau} - 1 < l_r$, since the minimum number of indel edges (as described above) are all insertion edges in that case. We compute the heuristic as follows:

**Definition 3** $$h\langle v, i, I\rangle = \begin{cases} 0 & \text{if } N^{\min}_g = 0 \\ N^{\min}_g \Delta_g & \text{if } d^{\max}_{v,\tau} - 1 < l_r \\ \Delta_o + N^{\min}_g \Delta_g & \text{otherwise} \end{cases}$$

Similarly, for a state $\langle v, i, D\rangle$, we would not have to incur the gap open cost again if $d^{\min}_{v,\tau} - 1 > l_r$, since the minimum number of indel edges are all deletion edges in that case. The heuristic is computed as follows:

**Definition 4** $$h\langle v, i, D\rangle = \begin{cases} 0 & \text{if } N^{\min}_g = 0 \\ N^{\min}_g \Delta_g & \text{if } d^{\min}_{v,\tau} - 1 > l_r \\ \Delta_o + N^{\min}_g \Delta_g & \text{otherwise} \end{cases}$$

#### Implementation details of superbubble-informed pruning

To ensure efficient detection of prunable states, POASTA keeps track of which bubble exits are reached during alignment. It stores for every node $v \in V$ the set of reached query positions in a B-tree set, which keeps the set of reached positions ordered. When testing an alignment state $\langle v, i\rangle$, from which it is possible to reach alignment states $\langle t, j^{\min}\rangle \ldots \langle t, j^{\max}\rangle$ (Main Text Figure 3), a B-tree enables quickly retrieving the query positions that previously reached the bubble exit in the range $[j^{\min}, j^{\max}]$.

POASTA considers the costs of implicitly opened indels, even when the superbubble is reached multiple times during alignment with varying costs (Supplemental Figure 3a). When a bubble exit is reached multiple times, POASTA tests whether the cost $C$ of an alignment state under test can improve over an implicitly opened insertion and an implicitly opened deletion from a different position (Supplemental Figure 3b).

A naive approach would be to test costs for the full range of positions $[j^{\min}, j^{\max}]$. POASTA, however, exploits the fact that the gap costs are linearly increasing with length. This means that POASTA only needs to check positions adjacent to previously reached exits, because those positions will have the lowest gap cost. For example, POASTA would only assess the gap costs of positions $j_2 - 1, j_2$, and $j_2 + 1$ in the example shown in Supplementary Figure 3b.

#### Construction of benchmark datasets

To construct our bacterial gene benchmark datasets, we first downloaded all bacterial "complete" genomes from NCBI RefSeq (40,188 genomes total; accessed July 2023). We used the accompanying gene annotations to extract the *dnaG*, *nusA*, *pgk*, *pyrG*, and *rpoB* gene sequences from each genome.

To create each individual benchmark set, we clustered gene sequences using single-linkage hierarchical clustering, as implemented in SciPy (Virtanen et al., 2020). Pairwise genetic distances were estimated using Mash (Ondov et al. (2016); $k = 15$; sketch size = 5,000), and were additionally used to deduplicate the sequence set, selecting one representative per set of identical sequences. We set the clustering threshold to 0.1, i.e., a new cluster would be formed if no neighbor could be found with a genetic distance < 0.1. This threshold is coarse enough to generate multiple genus and species-level clusters. We picked one or more clusters for each gene family as final datasets, each with at least 100 sequences, and varying the pairwise average nucleotide identities (ANI). Finally, each set was sorted by picking one "center" sequence with the smallest average Mash distance to all others and then ordering the remaining sequences in the set by the distance to the chosen "center" sequence, a strategy commonly applied before POA Gao et al. (2021).

#### Benchmark execution details

We ran POASTA with the following parameters: mismatch cost $\Delta_x = 4$, gap open cost $\Delta_o = 6$, and gap extend cost $\Delta_g = 2$, the same costs as used in the Wavefront Algorithm (WFA) (Marco-Sola et al., 2021). SPOA was configured to perform global alignment using the same cost model, and executed as follows:

```
spoa -l 1 -m 0 -n -4 -g -8 -e -2 -q 0 -c 0 -r 1
```

Tools were run in single-threaded mode on a `c2-standard-8` virtual machine on the Google Cloud Platform, with an Intel Cascade Lake CPU and with 32 GB of RAM.

#### Construction of **Mycobacterium tuberculosis** dataset

To construct the benchmark sets with *Mycobacterium tuberculosis* genomic sequences of 250, 500, and 1000 kbp in length, we downloaded all "complete" *M. tuberculosis* genomes available on NCBI RefSeq (370 total; accessed November 2023). To make all genomes colinear, we rotated and reoriented each genome such that each started with the gene *dnaA*, using the `fix-start` utility in Circlator (Hunt et al., 2015). Additionally, since inversions also break co-linearity, and POA poorly supports aligning large inversions, we excluded 29 genomes with more than 15% (≥ 660 kbp) of its genome inverted with respect to the canonical reference *M. tuberculosis* H37Rv, detected using MUMMER (Marçais et al., 2018).

We truncated genomes at specific genes to obtain sequences of the desired length. For the 250 kbp, 500 kbp, and 1 Mbp datasets, we used the genes *trmB*, *thiE*, and *gltA2* as cutoff points, respectively. We manually confirmed that these genes were located around the 250 kbp, 500 kbp, and 1 Mbp marks in each of the *dnaA* rotated and reoriented genomes. Finally, each dataset was sorted such that references were in ascending order of their Mash distance (Ondov et al., 2016) to H37Rv.

POASTA was executed with the same alignment cost model as described above, but on the larger `c2-standard-60` virtual machine on the Google Cloud Platform, which has 240 GB of RAM available.
